# Identification of a new Infectious Pancreatic Necrosis Virus (IPNV) isolate in Atlantic salmon (*Salmo salar* L.) that causes mortality in resistant fish

**DOI:** 10.1101/2021.05.23.445331

**Authors:** Borghild Hillestad, Stein Johannessen, Geir Olav Melingen, Hooman K. Moghadam

**Affiliations:** Benchmark Genetics Norway AS, Sandviksboder 3A, N-5035 Bergen, Norway; Benchmark Genetics, Sandviksboder 3A, N-5035 Bergen, Norway

**Keywords:** Infectious pancreatic necrosis (IPN), RNA sequencing, Genome assembly, Quantitative Trait Locus (QTL), Atlantic salmon

## Abstract

Infectious pancreatic necrosis (IPN) is an important viral disease of salmonids that can affect fish during various life cycles. In Atlantic salmon, selecting for genetically resistant animals against IPN has been one of the most highly praised success stories in the history of fish breeding. The findings that resistance against this disease has a significant genetic component, which is mainly controlled by variations in a single gene, has helped to reduce the IPN outbreaks over the past decade to a great extent. In this paper, we present the identification of a new isolate of the IPN virus, from a field outbreak, that had caused mortality, even in the genetically resistant animals. We recovered and assembled the full-length genome of this virus, following deep-sequencing of an infected tissue. The comparative sequence analysis revealed that for the critical amino acid motifs, previously found to be associated with the degree of virulence, the newly identified isolate is similar to the virus’s avirulent form. However, we detected a set of deduced amino acid residues, particularly in the hypervariable region of the polyprotein, that collectively are unique to this strain compared to all other reference sequences assessed in this study. We suggest that these mutations have likely equipped the virus with the capacity to escape the host defence mechanism more efficiently, even in the genetically deemed IPN resistant animals.

## Introduction

Infectious pancreatic necrosis (IPN) is one of the leading viral diseases of the Norwegian farmed Atlantic salmon (*Salmo salar* L.). The disease was first reported in 1941, following an outbreak in brook trout (*Salvelinus fontinalis* L.) in Canada (M’Gonigle, 1941). Further, the characterisation of the causative agent, the IPN virus (IPNV), was performed in 1960 (Wolf et al., 1960). In Norway, the virus was first isolated in 1975 from freshwater rainbow trout (*Oncorhynchus mykiss* L.) (Hastein and Krogsrud, 1976) and was designated as a notifiable disease from 1991 till 2008 (Sommerset et al., 2020). The virus can infect Atlantic salmon during all of its developmental stages, but the fish are especially susceptible as fry, during start-feeding, and post-smolts, soon after transfer to seawater (Roberts and Pearson, 2005; Sommerset et al., 2020). As suggested by the name, the pancreas is the primary target tissue of IPNV. However, the liver has also been shown to be one of the key organs affected by this virus (Ellis et al., 2010; Munang’Andu et al., 2013). In addition to significant economic losses, this disease is a concern for animal welfare, as survivors following infection can continue to infect naïve fish (Roberts and Pearson, 2005).

IPNV is a double-stranded RNA virus that belongs to the *Birnaviridae* family. A characteristic feature of the birnaviruses is their possession of a bi-segmented genome (i.e., A and B segments), contained within a non-enveloped, icosahedral capsid (Dobos and Roberts, 1983; Duncan et al., 1987; Duncan and Dobos, 1986). The smaller genomic segment, the B segment, is about 2.5 kb in size and consists of a single open reading frame (ORF), encoding an RNA-dependent, RNA polymerase VP1 (approximately 90 kDa) (Duncan et al., 1987; Duncan and Dobos, 1986). The A-segment is approximately 3 kb, and it contains two, partially overlapping ORFs (Duncan and Dobos, 1986). The first ORF, encodes VP5, a small, cysteine-rich, non-structural protein of about 17 kDa. A precursor polyprotein (NH2-preVP2-NS VP4 protease-VP3-COOH) of approximately 106 kDa constitutes the second and the larger ORF. The encoded VP4 protease cleavages the polyprotein into its two main components, the preVP2 and VP3. The preVP2 further matures to become VP2 and forms the outer capsid protein, containing the neutralizing epitopes and sites that facilitate cell attachment (Dopazo, 2020; Heppell et al., 1995b). The VP3 is the inner capsid protein (Duncan et al., 1987; Duncan and Dobos, 1986) but also in association with the VP1 it seems to be involved in viral packaging and replication (Tacken et al., 2002).

Based on cross-neutralizing tests, the aquatic birnaviruses are broadly divided into two main serogroups, A and B (Hill and Way, 1995). The serogroup A comprises of nine different serotypes, A_1_-A_9_. Most of the isolates from the USA fall within the A_1_ serotype (West Buxton; WB), the A_2_-A_5_ include mainly the European isolates (A_2_ (Spjarup; Sp), A_3_ (Abild; Ab), A_4_ (Hecht, He) and A_5_ (Te)), and A_6_-A_9_ are predominantly variants from Canada (A_6_ (Canada 1; C1), A_7_ (Canada 2; C2), A_8_ (Canada 3; C3) and A_9_ (Jasper)) (Blake et al., 2001; Hill and Way, 1995). The serogroup B consists of only one serotype (B1 (Tellinavirus; TV1)) that is non-pathogenic in fish. Based on the analysis of variations in the nucleotide and the deduced amino acids of the VP2, Blake et al. (2001) later proposed that aquatic birnaviruses constitute six major genogroups. These groups correspond relatively well with the geographical origins and serological classifications previously suggested. According to this classification, WB and the Jasper strains constitute genogroup 1, the A_3_ serotype forms genogroup 2, C1 and Te make genogroup 3, the two Canadian strains, C2 and C3 form genogroup 4, all European isolates of serotype A_2_ cluster into genogroup 5 and the He strain constitutes genogroup 6 (Blake et al., 2001). A seventh genogroup was also suggested based on isolates recovered from Japanese aquatic sources (Nishizawa et al., 2005).

Despite extensive vaccination efforts, disinfection and targeted breeding programs, IPN outbreaks remains a concern in farmed Atlantic salmon. Related mortalities can be substantial and in part, can be a function of the genetic makeup of the host (Guy et al., 2006; Houston et al., 2008; Moen et al., 2009), management and environmental factors (Jarp et al., 1995; Sundh et al., 2009), and the strain and specific genetic variations carried by the virus (Skjesol et al., 2011; Song et al., 2005). There are ample examples of isolates, belonging to the same serotype, to cause different degrees of virulence in the host (e.g., Bruslind and Reno, 2000; Santi et al., 2004; Shivappa et al., 2004), and indeed, there have been many studies, investigating the molecular basis of IPNV virulence in Atlantic salmon (e.g., Mutoloki et al., 2016; Santi et al., 2004, 2005; Skjesol et al., 2011; Song et al., 2005). Although the genetic details of such processes are not yet fully understood, most studies speculated, and chiefly investigated, the genetic variations in segment A concerning the degree of virulence in IPNV. In particular, specific amino acid motifs in the VP2 of the polyprotein have been suggested to be most prevalent and in association with the virulent forms of the pathogen (Bruslind and Reno, 2000; Munang’andu et al., 2016; Mutoloki et al., 2016; Santi et al., 2004; Shivappa et al., 2004; Song et al., 2005). Variations in the amino acid residues in positions 217, 221 and 247 of the VP2 have been of particular focus (Dopazo, 2020; Julin et al., 2013; Munang’andu et al., 2016; Mutoloki et al., 2016; Santi et al., 2004; Song et al., 2005). Further, assessment of the recombinant viral strains, have indicated that the residue 217 of the VP2 might be the most crucial amino acid that can result in the loss or gain of virulence (e.g., Song et al., 2005).

Despite concerns and economic losses due to infection by IPNV, increased resistance against this virus through selective breeding is among the most remarkable success stories in the history of aquaculture and salmon breeding in particular. The initial attempt to estimate the genetic parameters controlling this trait, which was based on mortality in the field and controlled challenge testing, had shown that resistance to IPNV has a significant and heritable genetic component (*h^2^* of 0.17-0.62) (Kjøglum et al., 2008; Wetten et al., 2007). However, the breakthrough came with the finding that resistance to IPNV is mainly controlled by variations in a single quantitative trait locus (QTL) on chromosome 26 (Houston et al., 2008, 2010; Moen et al., 2009). Subsequent work identified mutations in the epithelial cadherin gene (*cdh1*) as the causative genetic variation for IPNV resistance in Atlantic salmon (Moen et al., 2015). Following the inclusion of this QTL into the breeding programs in Norway, the industry witnessed a sharp decline in the number of IPN outbreaks throughout the country, dropping from 223 to only 19 reported cases from 2009 to 2019 (Sommerset et al., 2020).

This paper reports the identification and whole-genome sequence assembly and analysis of a new IPNV variant. The isolate was recovered from a field outbreak in West of Norway following reported mortalities, due to an IPN infection, among QTL homozygous or heterozygous IPN resistant fish. We describe the amino acid motifs that distinguish this isolate from other phylogenetically related variants and suggest that these mutations are likely to play an essential role in this isolate’s pathogenicity.

## Materials and Methods

### Field outbreak and sample collection

In May of 2019, there was an IPN field outbreak in a Western region of Norway. The affected fish were Atlantic salmon of the SalmoBreed strain, delivered as IPN QTL, all favourable homozygous or heterozygous for the QTL locus (Houston et al., 2008; Moen et al., 2009). Fish were at the post-smolt stage with an approximate average weight of about 700 – 800 gr at the time of the outbreak. The reported mortality reached about 10%, a high percentage for field outbreak among QTL carrying fish. The IPNV infection was identified following an inspection by the veterinary authorities. The diagnosis was later validated by detection of the VP2 segment of the viral genome using polymerase chain reaction (qRT-PCR) of the infected head-kidney and through histopathological and immunohistochemical examinations of the liver and pancreatic tissues (Figure 1A and B), all performed at the Fish Vet Group Norway (http://fishvetgroup.no) (Supplementary Table 1). Tissues were fixed in a 4% formalin solution (4% formalin, 0.08M sodium phosphate, pH 7.0), processed using Thermo Scientific Excelsior^®^ and embedded in Histowax with the aid of Tissue – Tek^®^, TEC 5 (Sakura) embedding system. The embedded tissues were then sectioned at 1.5-2 μm using a Leica RM 2255 Microtome and stained with Hematoxylin-Eosin (HE). The stained sections were scanned in an Aperio ScanScope AT Turbo slide scanner and read using Aperio Image Scope (Leica Biosystems). The scans were examined and scored in a blind fashion, i.e., without information about the animal’s history. A clip from the adipose fin was also sampled and stored in 95% ethanol for DNA extraction and genotyping of ten fish, five moribund and five dead. We genotyped these fish for the genetic markers used to assign individuals as QTL carrier (Houston et al., 2008; Moen et al., 2009, 2015), and confirmed all animals carry IPN QTL.

**Figure 1.**
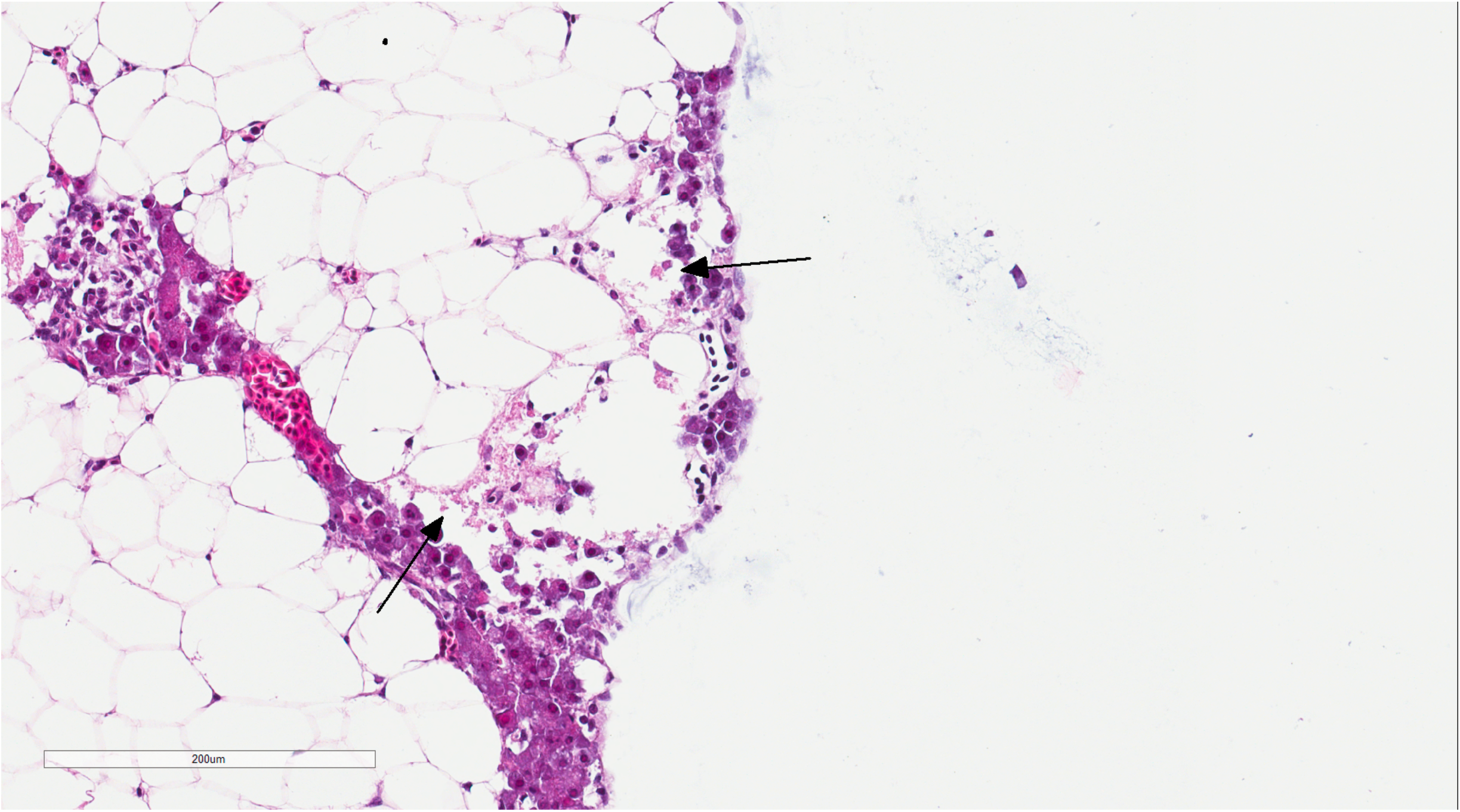
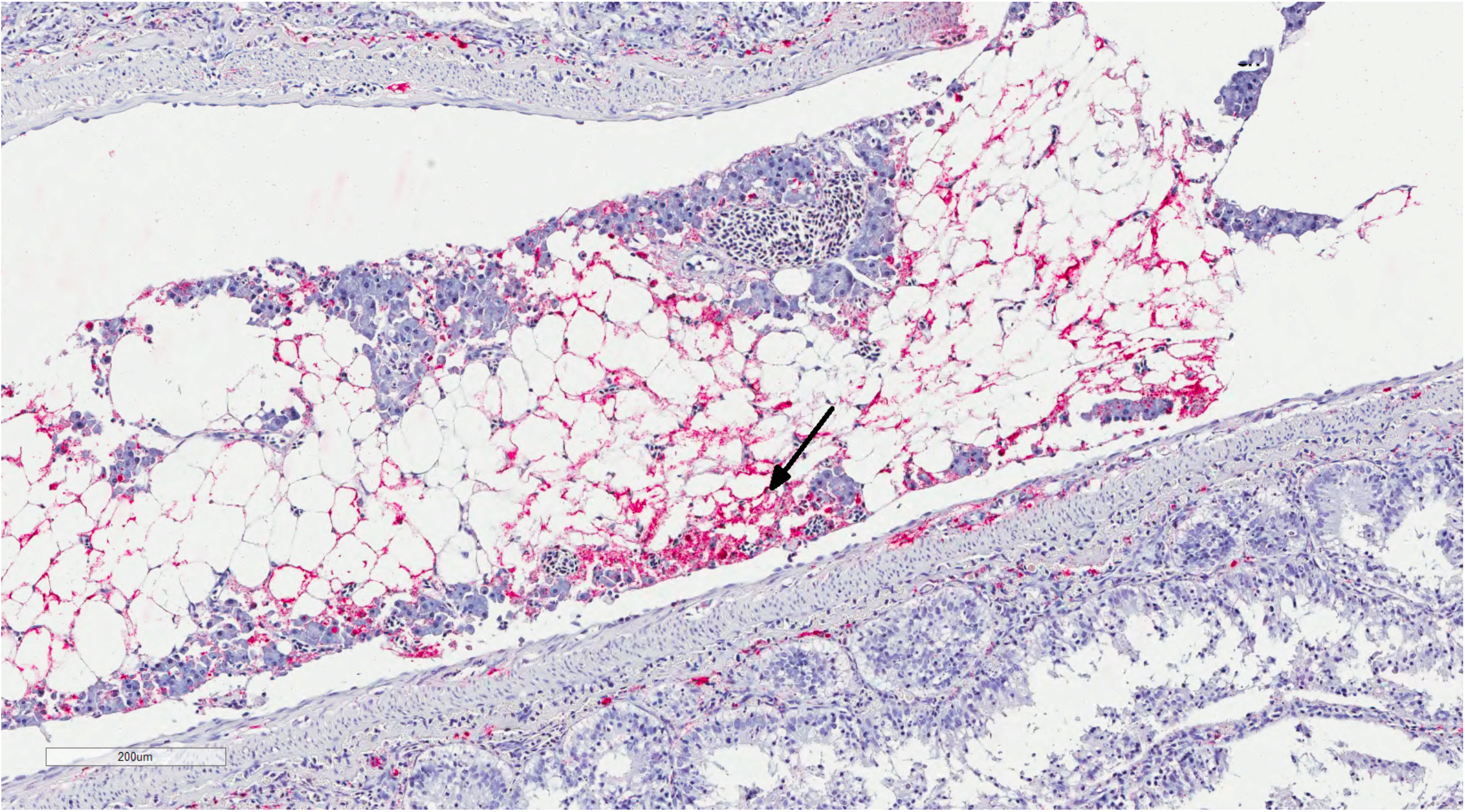
A. Histology section of the pancreatic tissue infected with the infectious pancreatic necrosis virus (IPNV). Arrows indicate acute necrosis of pancreatic acinar cells. B. Immunohistochemical staining of the pancreas tissue using antibodies specific for IPNV. Red colours show cells infected with this virus (arrow).

### Transcriptome sequencing and viral genome assembly

The head-kidney was collected from six infected fish (http://fishvetgroup.no/pcr-2/#1452163253406-d81f6dce-bdf9), all at a moribund state, and stored in RNALater^®^ (Ambion). Total RNA extraction was performed using the RNeasy^®^ Plus mini kit (Qiagen). Nanodrop ND-1000 Spectrophotometer (NanoDrop Technologies) was used to assess the concentration and purity of the extracted genetic materials. The Norwegian Sequencing Centre performed library construction and sequencing the total RNA on an Illumina HiSeq 4000 platform as paired-end (PE) 150 bp reads.

We first cleaned the sequence data by removing low-quality nucleotides and sequencing adapters using Trimmomatic (v.0.36) (Bolger et al., 2014). The ribosomal RNA (rRNA) was then identified through a similarity search against the SILVA rRNA database (Quast et al., 2013) and excluded from subsequent analysis. The host-specific transcriptome was detected by aligning the remaining reads against the Atlantic salmon reference genome assembly (ICSASG_v2) (Lien et al., 2016) using TopHat (v.2.0.13) (Trapnell et al., 2009, 2012). The un-aligned reads were then blasted against the IPNV reference assemblies (ASM397166v1, ASM397170v1, ASM397174v1, ASM397168v1, ASM397172V1 and ViralMultiSegProj15024) by setting “task” as “megablast”, “word size” to 7 and “expectation value threshold” to 1.00e^-07^. All the reads that aligned to the IPNV reference sequences were extracted, pooled into a single sequence datafile per animal, and then fed into Trinity (v.2.11.0) (Grabherr et al., 2011) with the default parameter settings to construct assemblies.

### Comparative genomics and phylogenetic analysis

We compared the assembled viral genomes to one another (Supplementary Tables 1 and 2), and against various reference isolates, with different degrees of virulence (Dopazo, 2020; Julin et al., 2013; Munang’andu et al., 2016; Mutoloki et al., 2016; Santi et al., 2004; Song et al., 2005) (Table 1) or strains representing different genogroups, as previously suggested (Blake et al., 2001) (Table 2). Further, we assessed the deduced amino acids of these isolates to any IPNV sequence in the NCBI database with a complete polyprotein or VP1 information (Supplementary Files 1 and 2). The sequences included 90 isolates with the full coding sequence for the polyprotein and 120 for the VP1. Both, the nucleotide and amino acid alignments were performed using ClustalX (Larkin et al., 2007) implemented in Unipro UGENE (v36.0) (Okonechnikov et al., 2012) or Muscle (Edgar, 2004) implemented in Seqotron (v1.0.1) (Fourment and Holmes, 2016). Unipro UGENE and Mega X (v10.1.7) (Kumar et al., 2018) were also used to assess sequence similarity, calculate pair-wise distances for nucleotide and amino acid data, and to construct phylogenetic associations using the neighbour-joining (NJ) method. To assess the confidence of node assignments, we performed 1000 bootstrap replications.

**Table 1.** Amino acid sequences in segment A of the newly identified IPNV isolate compared with previously reported key residues suggested being correlated with viral virulence (Santi et al., 2004).

| Accession | Isolate | 217 | 221<sup>a</sup> | 247 | 252 | 500 | Virulence |
| --- | --- | --- | --- | --- | --- | --- | --- |
| AY374435.1 | NVI-001 | T | A/T | T | V | Y | avirulent |
| AY379742.1 | NVI-016 | P | T | A | N | Y | avirulent |
| AY379744.1 | NVI-010 | P | T | A | N | H | moderate |
| AY379735.1 | NVI-011 | T | T/A | A | V | H | virulent |
| AY379738.1 | NVI-013 | T | A/T | T | V | Y | virulent |
| AY379740.1 | NVI-015 | T | A/T | T | V | Y | virulent |
| AY379736.1 | NVI-020 | T | T/A | A | V | H | virulent |
| AY379737.1 | NVI-023 | T | A/T | T | V | Y | virulent |
| - | BG | P | T | A | D | Y | - |

<sup>a</sup> Double peak on the chromatograms after two cell culture passages. The dominating amino acid is indicated first.

**Table 2.** Amino acid residues and their positions in the polyprotein region of the IPNV isolates reported in this study that are either exclusively unique or occur with a very low frequency in other strains.

| Num. | Accession | ID | 245 | 248 | 252 | 255 | 257 | 278 | 282 | 285 | 321 | 585 |
| --- | --- | --- | --- | --- | --- | --- | --- | --- | --- | --- | --- | --- |
| 1 | - | BG | G | R | D | T | H | A | T | H | D | G |
| 2 | AJ829474 | Sp_975/99 | S | E | I | K | D | V | N | Y | G | D |
| 3 | AY379735 | Sp_NVI-011 | S | E | V | K | D | V | N | Y | G | D |
| 4 | AY379738 | Sp_NVI-013 | S | E | V | K | D | V | N | Y | G | D |
| 5 | AY379740 | Sp_NVI-015 | S | E | V | K | D | V | N | Y | G | D |
| 6 | AY379736 | Sp_NVI-020 | S | E | V | K | D | V | N | Y | G | D |
| 7 | AY379737 | Sp_NVI-023 | S | E | V | K | D | V | N | Y | G | D |
| 8 | AY379744 | Sp_NVI-010 | S | E | N | K | D | V | N | Y | G | D |
| 9 | AY374435 | Sp_NVI-001 | S | E | V | K | D | V | N | Y | G | D |
| 10 | AY379742 | Sp_NVI-016 | S | E | N | K | D | V | D | Y | G | D |
| 11 | AF343573 | Buhl | Q | A | N | V | D | T | V | H | A | D |
| 12 | AF342729 | Ab | S | N | A | K | D | V | A | Y | A | D |
| 13 | AF342730 | He | N | Q | N | K | D | V | E | Y | G | D |
| 14 | AF342731 | Te | R | A | A | K | D | V | A | F | T | D |
| 15 | AF342732 | C1 | S | R | T | R | D | V | A | F | T | D |
| 16 | AF342733 | C2 | S | T | A | T | E | A | K | H | G | D |
| 17 | AF342734 | C3 | S | N | A | T | K | A | T | H | A | D |
| 18 | AF342735 | Jasper | Q | A | N | R | D | A | A | H | G | D |
| 19 | AF342727 | WB | Q | A | N | R | D | T | A | Y | G | D |

## Results and Discussion

### Transcriptome sequencing and genome assembly

This paper describes the genomic features of IPNV isolates, responsible for a field outbreak in an Atlantic salmon population. The post-mortem examination identified IPN infection as the chief cause of mortality among all fish investigated. The histopathological and immunohistochemical examinations (Figure 1A and B and Supplementary Table 1) and the viral genetic material detected through qRT-PCR amplification (Supplementary Table 1) further confirmed earlier diagnosis. We found that the genotypes of all animals examined in this study were consistent with an IPN resistant animal (Houston et al., 2008; Moen et al., 2009, 2015). Further, assessment of the transcriptome sequence data confirmed that all fish carried at least one favourable copy of the causal mutation in the *cdh1* gene, conferring resistance against IPNV (Moen et al., 2015). The causal mutation, which is in the form of a single nucleotide polymorphic (SNP) variation, is located in the protein-coding sequence of *cdh1* (ssa26:15192533; C/T) and can change Proline (P) to Serine (S), with the former amino acid found to be associated with the IPNV resistance (Moen et al., 2015). It is expected that in a heterozygote fish, the favourable allele exhibit close to complete dominant effect in response to the virus (Moen et al., 2015). Among the animals investigated in this study, three fish were homozygous for the favourable allele (C/C), and the other three were heterozygous (C/T). The extracted RNA was sequenced to high depth, with an average of 60 million PE reads per animal. Following the rRNA’s exclusion, more than 45 million PE reads per fish remained for the assemblage of the host’s viral genome and profiling gene expression. Together, the two segments of the IPNV genome consist of about 5800 nucleotides. On average, we obtained 56-fold coverage of the viral genome from each of the animals in our sequence data. Here, we refer to the assembled viral genomes as BG_x_y, where x denotes the animal’s id (1-6) and y refers to the viral genome’s specific segment (e.g., VP1 or polyprotein).

### Comparative sequence analysis of the polyprotein

Comparative genomic analysis between the six assembled genomes revealed at least two different, closely linked variants, caused the field outbreak in the population investigated (Figure 2A and B; Supplementary Table 2A and B). The nucleotide sequences of the polyprotein are identical in four of the assemblies (BG_1_polyprot to BG_4_polyprot) and contain nine polymorphic sites compared to the other two assembled sequences (i.e., BG_5_polyprot and BG_6_polyprot). Two of the polymorphisms are nonsynonymous, changing the amino acid residues in positions 47 (aspartate (D) to glutamate (E)) and 717 (lysine (K) to glutamine (Q)).

**Figure 2.**
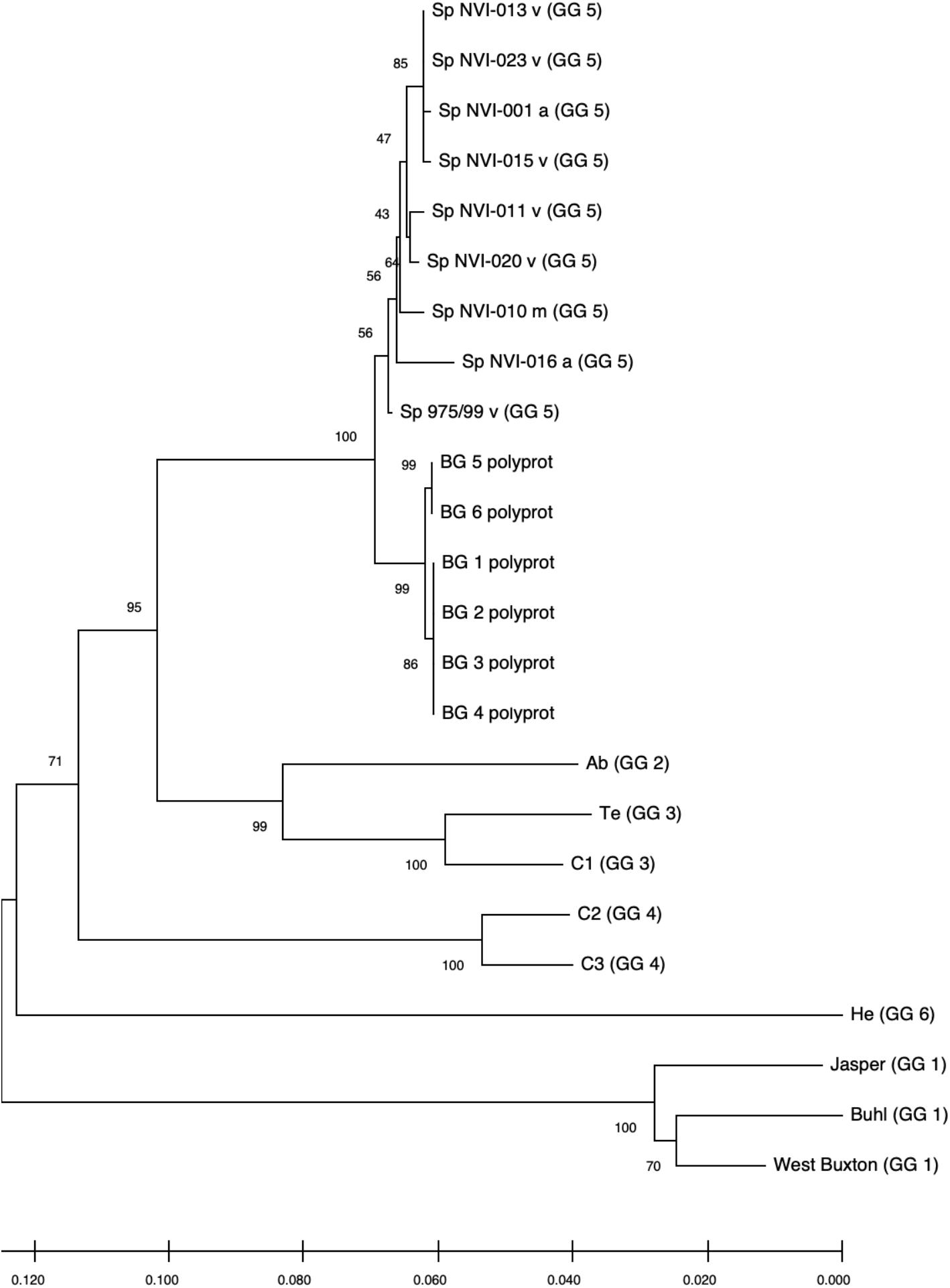
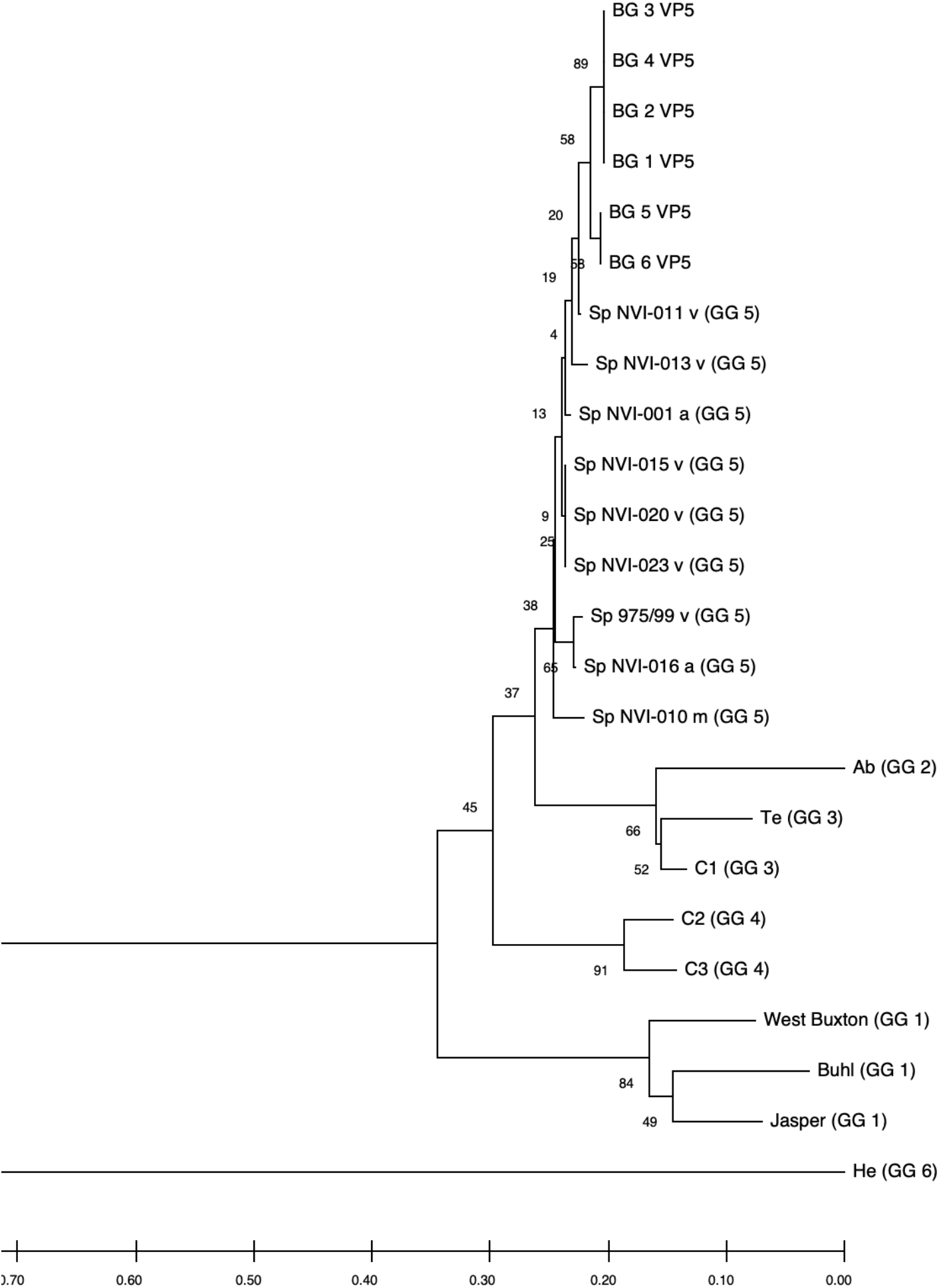
Neighbour-Joining tree, showing the phylogenetic relationship between the A. Polyprotein and B. VP5’s deduced amino acid data from different IPNV reference strains, and the assemblies reported in this study. The accession ids of the sequences are presented in Table 2. The associated genogroups are in parentheses. BG 1 to 6 represent isolates identified in the current work. The confidence of association between sequences was estimated using bootstrap testing (1000 replicates) and is shown next to the branches. The Maximum Composite Likelihood method (Tamura K et al., 2004) was used to estimate the isolates’ evolutionary distances.

In a study, aiming to trace the IPNV infection signatures from hatcheries to the sea (Kristoffersen et al., 2018), the authors have recovered a partial fragment of the polyprotein from an isolate (accession number: MH562009), with very similar nucleotide sequence to the ones reported here. Recently, in controlled infection challenge testing, the mortality induced by this isolate among full-sib families was compared with a standard, virulent form of the virus (AY379740.1). While the infection with the latter variant resulted in approximately 30% mortality, mainly among the non-QTL fish, the former isolate caused more than 75% mortality, treating the QTL and non-QTL fish alike (Ørpetveit et al. in preparation; https://www.vetinst.no/nyheter/endringer-i-ipn-viruset-gjor-fisken-mer-utsatt-for-sykdom, https://www.fishfarmingexpert.com/article/changes-in-ipn-virus-make-salmon-more-susceptible, https://salmobreed.no/articles/benchmark-genetics-intensiverer-forskningen-pa-nytt-ipn-virus). Comparison between the nucleotide sequences of this isolate with our assembled genomes shows divergence ranging from 0.003-0.004 (Supplementary Figure 1A). However, this isolate has deduced amino acid sequences identical to the translated sequences from our first four assemblies (Supplementary Figure 1B), indicating that this variant is likely to be the ancestral form and the other variants (i.e., BG_5_VP2 and BG_6_VP2) are more recently derived. One of the main characteristics of RNA viruses is their high mutation rates, mainly attributed to their lack of an effective proofreading activity. Therefore, finding different sequence variants from the same outbreak might not be surprising, but it indeed is important, as it will allow one to follow the trajectory of pathogen evolution through time and space.

Comparative assessment of the polyprotein nucleotide (data not shown) as well as the amino acid sequence data with multiple reference strains (Figure 2A) shows our assemblies are most closely related to the Sp serotype (genogroup 5). However, they also constitute a distinct group within this serotype compared to Sp isolates assessed in this study (Figure 2A; Supplementary Figure 2A). Several investigations have previously reported variations in some amino acid residues, particularly in the VP2 of the polyprotein, to be associated with the degree of the IPNV virulence (Bruslind and Reno, 2000; Munang’andu et al., 2016; Mutoloki et al., 2016; Santi et al., 2004; Shivappa et al., 2004; Song et al., 2005). Interestingly, the amino acid motifs found in the six assembled genomes in this study are mainly consistent with the virus’s avirulent form (Table 1) (Santi et al., 2004; Shivappa et al., 2004; Song et al., 2005). For instance, for the most critical positions associated with the virulence, i.e., 217, 221 and 247 of the VP2, the amino acid residues of our assembled genomes are proline (P), threonine (T) and alanine (A) (Table 1; Supplementary File 1), while it is expected that in the virulent form, the corresponding residues to be T, A and T, respectively (Mutoloki et al., 2016; Santi et al., 2004; Song et al., 2005). On the other hand, we identified ten amino acid residues in the assembled genomes’ polyprotein, that either exclusively or with a disproportionate frequency are found in the reported isolates (Table 2; Supplementary File 1). Nine of these sites are within the VP2, and one is located in the VP4. In the VP2, these residues are all in proximity to one another, encompassing the hypervariable region (Blake et al., 2001; Heppell et al., 1995b). Specifically, these residues are located in positions 245, 248, 252, 255, 257, 278, 282, 285, 321 (Table 2; Supplementary File 1). This data provides additional support for the previously suggested notion that the amino acid motifs in the VP2 capsid protein hypervariable region, might play an essential role in the degree of the virulence displayed between different IPNV isolates (Bruslind and Reno, 2000; Skjesol et al., 2011). It is also likely that this specific constellation of the amino acid residues reported in these isolates has helped the virus to escape the host defence barrier, despite fish being regarded as resistant to IPNV. In the VP4, the deduced amino acid residue, unique to the assembled isolates (Table 2; Supplementary File 1), is located in position 585. The amino acid substitution is a glycine (G) residue, compared to the other sequences investigated in this study, where they carry an aspartate (D) (Table 2).

### Comparative sequence analysis of VP5

All assemblies contain the VP5, the partial overlapping, non-structural protein, of segment A. Similar to the polyprotein, the phylogenetic analysis of the VP5, grouped the assemblies into two distinct groups and further clustered them with the isolates from the Sp serotype (Figure 2B). The two assembled groups differ in only one amino acid residue at position 90, with the first four assemblies carrying an arginine (R) while the two other assemblies are having a glycine (G) (Supplementary File 3). Sequence analysis of the deduced amino acids of VP5 did not reveal any unique residue in the assembled isolates compared to all other sequences. Comparing to the other Sp serotypes, however, the amino acid in position 98 differs between the isolates (Supplementary File 3). At this position, all the investigated Sp isolates carry an R while in the assemblies, the residue is tryptophan (W). It has been shown that the VP5’s protein is generally produced during the initial stages of replication (Heppell et al., 1995a), and at least in the Ab serotype, it seems that this protein to be involved in anti-apoptotic functions (Hong and Wu, 2002). However, this product’s importance in viral growth or virulence has so far remained ambiguous (e.g., Julin et al. 2013; Dopazo 2020).

### Comparative sequence analysis of VP1

The phylogenetic analysis of VP1 clustered the assemblies into three distinct clades (Supplementary Figure 2B) with the amino acid variations in positc’s 119 (threonine (T) to methionine (M)) and 701 (T to alanine (A) and serine (S)) (Supplementary File 2; Supplementary Table 3A). The nucleotide sequence comparisons, shows a greater rate of divergence in the VP1 of the assemblies compared to those of the polyprotein, suggesting that the B segment of the genome has a higher rate of nucleotide mutation (Supplementary Tables 1 and 2). Comparing the deduced amino acid sequences of the assembled genomes to the VP1 section of 120 IPNV sequences, further revealed only a single, almost unique residue in our isolates (Supplementary File 2). The amino acid residue in position 656 of the assembled isolates is glutamate (E), while in the majority of other strains, the amino acid is either aspartate (D) or alanine (A). The only other isolates, carrying an E in this position, are two sequences recovered from Atlantic salmon (KY548520) and Brown trout (*Salmo trutta* L.) (KY548519) in Finland following outbreaks (Holopainen et al., 2017). However, it should be noted that so far, the role of VP1 in determining the degree of IPNV virulence has remained ambiguous. For example, in a study conducted by Song et al. (2005), through construction and comparative assessment of chimeric IPNV strains, the authors suggested that VP1 is not involved in this pathogen’s virulence. On the other hand, the data reported by Shivappa et al. (2004), is suggestive that the amino acids on the B segment of the genome in combination with specific residues in VP2, might help to modulate the magnitude of virulence. Work in the infectious bursal disease virus (IBDV) has also shown that variations in the amino acid residues of the VP1 can change the kinetics of viral replication and influence the virus’s virulence (Liu and Vakharia, 2004). VP1 interacts with VP3 to form VP1-VP3 complexes, and as such, can affect various aspects of the virus biology, including its replication efficiency (Pedersen et al., 2007; Tacken et al., 2002).

### Concluding remarks

In light of the findings reported in this study, a key point worth noting is the importance of understanding the genetic interactions between the host and the pathogen in determining an infection’s outcome. As a general trend, research studies tend to mainly focus on either the host or the pathogen’s genetic polymorphisms while investigating the genetic basis of disease-related phenotypic variations. While the merits of such a targeted approach cannot be disputed, it is becoming increasingly important also to be aware of the detailed genetic and genomic variations and interactions between the host and the microbe. In the case of monogenic infectious diseases, it is relatively trivial to speculate whether a new form of a pathogen interacts in fundamentally different ways with the host’s defence mechanism. For instance, considering that resistance to IPNV in Atlantic salmon is mainly determined by variations in a single gene, one might expect that the mutations acquired by the virus strain reported in this paper might have offered an adaptive advantage, assisting the pathogen to establish successful infection, irrespective of the variations in the *cdh1* gene. Of course, pinpointing the causative mutations and understanding and validating the molecular basis of such adaptive variations is a substantial undertaking by itself. The task of identifying and untangling the dynamics of how genotypic variations in the host and the pathogen influence one another and affect the outcome of an infection becomes more challenging while investigating diseases with polygenetic nature. This point becomes even more critical, considering our rapidly changing environments that can facilitate the transmission dynamics and the geographical spread of the pathogens (Reid et al., 2019). For instance, changes in water temperature, pH and oxygen have been linked to saprolegniasis in the Indian major carps (*Catla catla, Labeo rohita* and *Chirrinus mrigal*) (Das et al., 2012) and to white spot syndrome virus in prawns (*Penaeus monodon* and *Fenneropenaeus indicus*) (Selvam et al., 2012). Fortunately, advancements in technologies such as sequencing, genotyping, computational analysis and molecular biology can now provide us with many necessary tools, if we are set to understand the dynamics of the interactions between the host and the microbe at their detailed molecular levels.

## Data availability

All datasets generated for this study are included in the article (Supplementary Data) and have also been deposited in GenBank under the accession numbers MW496366, MW496367, MW496368, MW496369, MW496370, MW496371, MW496372, MW496373, MW496374, MW496375, MW496376 and MW496377.

## Acknowledgements

We would like to thank the Fish Vet Group Norway and especially Dr Marianne Kraugerud and Mr Simon Rey for all the laboratory work and the Norwegian Sequencing Centre for the excellent work and guidance.

## Contributions

G.M., S.J. and B.H. arranged for tissue sampling, shipment and all laboratory and sequencing procedures. H.M. performed analyses of the sequence data. B.H. and H.M. drafted the manuscript. All authors contributed to the development of the paper, interpretation of the results and improvement of the manuscript.

## Supplementary Figure Legends

**Supplementary Figure 1.** Neighbour-Joining trees, showing the phylogenetic relationships between partial polyprotein nucleotide (A) and the amino acid (B) sequences of the reported assemblies with an isolate (MH562009) previously recovered from a field outbreak (Kristoffersen et al., 2018). BG 1 to 6 represent assembled sequences in the current study. The distances were computed using the Maximum Composite Likelihood method, for nucleotides and Poisson for the amino acid data. The rate variation among sites was modelled with a gamma distribution (shape parameter = 1). There were a total of 1513 nucleotides and 504 amino acids in the final dataset. Evolutionary analyses were conducted in MEGA X.

**Supplementary Figure 2.** Neighbour-Joining trees, showing the phylogenetic relationships between the assembled polyprotein (A) and VP1 (B) segments of the IPNV genomes for the six samples sequenced in this study. The trees are constructed based on the amino acid sequence information. The confidence of association, estimated using bootstrap testing (1000 replicates), is shown next to the branches. There were 2919 and 845 positions in the final datasets for the polyprotein and VP1 respectively. Analyses were conducted in MEGA X.

**Supplementary Table 1.** Overview of the laboratory results, confirming the initial diagnosis of the IPNV infection and QTL status of the fish.

| Num. | tissue | QTL | qRT-PCR Ct value | histology | Immunohistochemistry |
| --- | --- | --- | --- | --- | --- |
| 1 | adipose fin | Yes | Not tested | Not tested | Not tested |
| 2 | adipose fin | Yes | Not tested | Not tested | Not tested |
| 3 | adipose fin | Yes | Not tested | Not tested | Not tested |
| 4 | adipose fin | Yes | Not tested | Not tested | Not tested |
| 5 | adipose fin | Yes | Not tested | Not tested | Not tested |
| 6 | adipose fin | Yes | Not tested | Not tested | Not tested |
| 7 | adipose fin | Yes | Not tested | Not tested | Not tested |
| 8 | adipose fin | Yes | Not tested | Not tested | Not tested |
| 9 | adipose fin | Yes | Not tested | Not tested | Not tested |
| 10 | adipose fin | Yes | Not tested | Not tested | Not tested |
| 11 | kidney | Not tested | 20.32 | Not tested | Not tested |
| 12 | kidney | Not tested | 19.79 | Not tested | Not tested |
| 13 | kidney | Not tested | 18.93 | Not tested | Not tested |
| 14 | kidney | Not tested | 18.92 | Not tested | Not tested |
| 15 | kidney | Not tested | 20.77 | Not tested | Not tested |
| 16 | kidney | Not tested | 19.32 | Not tested | Not tested |
| 17 | multiple | Not tested | Not tested | Multifocal necrosis in the exocrine pancreas | positive |
| 18 | multiple | Not tested | Not tested | Multifocal necrosis in exocrine pancreas and multifocal bleeding in abdominal fat | positive |
| 19 | multiple | Not tested | Not tested | Multifocal necrosis in exocrine pancreas and focal, extensive necrosis in the liver | positive |

**Supplementary Table 2.** Estimates of evolutionary divergence rate between the amino acid (A) and nucleotide (B) sequences of the polyprotein of the assembled genomes. The actual number of differences are provided in the parentheses. The rate variation among sites was modelled with a gamma distribution (shape parameter = 1).

| **A** | **BG_1_poly** | **BG_2_poly** | **BG_3_poly** | **BG_4_poly** | **BG_5_poly** | **BG_6_poly** |
| --- | --- | --- | --- | --- | --- | --- |
| **BG_1_poly** | 0 (0) | 0 (0) | 0 (0) | 0 (0) | 0.00206 (2) | 0.00206 (2) |
| **BG_2_poly** |  | 0 (0) | 0 (0) | 0 (0) | 0.00206 (2) | 0.00206 (2) |
| **BG_3_poly** |  |  | 0 (0) | 0 (0) | 0.00206 (2) | 0.00206 (2) |
| **BG_4_poly** |  |  |  | 0 (0) | 0.00206 (2) | 0.00206 (2) |
| **BG_5_poly** |  |  |  |  | 0 (0) | 0 (0) |
| **BG_6_poly** |  |  |  |  |  | 0 (0) |

| **B** | **BG_1_poly** | **BG_2_poly** | **BG_3_poly** | **BG_4_poly** | **BG_5_poly** | **BG_6_poly** |
| --- | --- | --- | --- | --- | --- | --- |
| **BG_1_poly** | 0 (0) | 0 (0) | 0 (0) | 0 (0) | 0.00308 (9) | 0.00308 (9) |
| **BG_2_poly** |  | 0 (0) | 0 (0) | 0 (0) | 0.00308 (9) | 0.00308 (9) |
| **BG_3_poly** |  |  | 0 (0) | 0 (0) | 0.00308 (9) | 0.00308 (9) |
| **BG_4_poly** |  |  |  | 0 (0) | 0.00308 (9) | 0.00308 (9) |
| **BG_5_poly** |  |  |  |  | 0 (0) | 0 (0) |
| **BG_6_poly** |  |  |  |  |  | 0 (0) |

**Supplementary Table 3.** Estimates of evolutionary divergence rate between the amino acid (A) and nucleotide (B) sequences of the VP1 of the assembled genomes. The actual number of differences are provided in the parentheses. The rate variation among sites was modelled with a gamma distribution (shape parameter = 1).

| **A** | **BG_1_VP1** | **BG_2_VP1** | **BG_3_VP1** | **BG_4_VP1** | **BG_5_VP1** | **BG_6_VP1** |
| --- | --- | --- | --- | --- | --- | --- |
| **BG_1_VP1** | 0 (0) | 0.00118 (1) | 0 (0) | 0.00118 (1) | 0.00237 (2) | 0.00237 (2) |
| **BG_2_VP1** |  | 0 (0) | 0.00118 (1) | 0 (0) | 0.00237 (2) | 0.00237 (2) |
| **BG_3_VP1** |  |  | 0 (0) | 0.00118 (1) | 0.00237 (2) | 0.00237 (2) |
| **BG_4_VP1** |  |  |  | 0 (0) | 0.00237 (2) | 0.00237 (2) |
| **BG_5_VP1** |  |  |  |  | 0 (0) | 0 (0) |
| **BG_6_VP1** |  |  |  |  |  | 0 (0) |

| **B** | **BG_1_VP1** | **BG_2_VP1** | **BG_3_VP1** | **BG_4_VP1** | **BG_5_VP1** | **BG_6_VP1** |
| --- | --- | --- | --- | --- | --- | --- |
| **BG_1_VP1** | 0 (0) | 0.00158 (4) | 0.000395 (1) | 0.00158 (4) | 0.00592 (15) | 0.00553 (14) |
| **BG_2_VP1** |  | 0 (0) | 0.00118 (3) | 0 (0) | 0.00474 (12) | 0.00434 (11) |
| **BG_3_VP1** |  |  | 0 (0) | 0.00118 (3) | 0.00553 (14) | 0.00513 (13) |
| **BG_4_VP1** |  |  |  | 0 (0) | 0.00474 (12) | 0.00434 (11) |
| **BG_5_VP1** |  |  |  |  | 0 (0) | 0.000395 (1) |
| **BG_6_VP1** |  |  |  |  |  | 0 (0) |

## Supplementary Files

**Supplementary File 1.** Alignment of the deduced amino acid sequence data from the polyprotein section of the IPNV. BG 1 to 6 represent the six assemblies from the current study.

**Supplementary File 2.** Alignment of the deduced amino acid sequence data from the VP1 section of the IPNV. BG 1 to 6 represent the six assemblies from the current study.

**Supplementary File 3.** Alignment of the deduced amino acid sequence data from the VP5 section of the IPNV. BG 1 to 6 represent the six assemblies from the current study.

