## Supplementary figures and images for "Identification of a new Infectious Pancreatic Necrosis Virus (IPNV) isolate in Atlantic salmon (*Salmo salar* L.) that causes mortality in resistant fish"

### Data Sheet 4.PDF

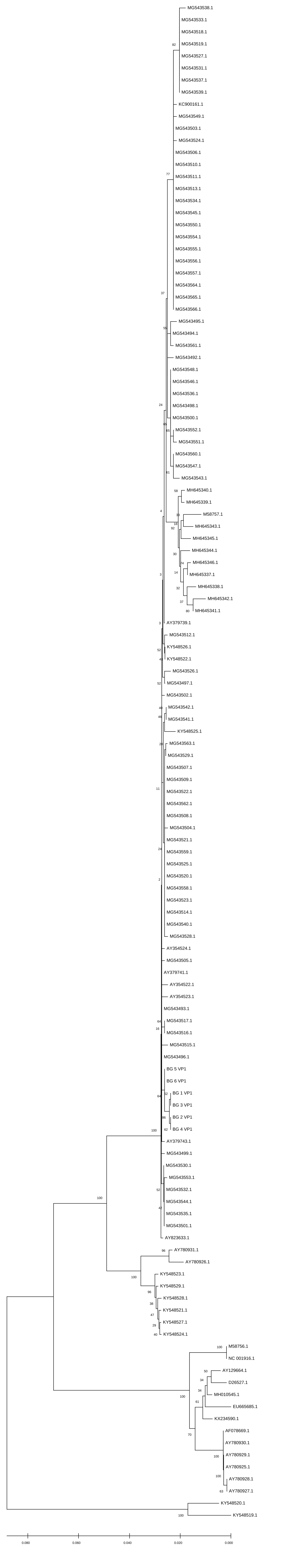

### Data Sheet 5.PDF

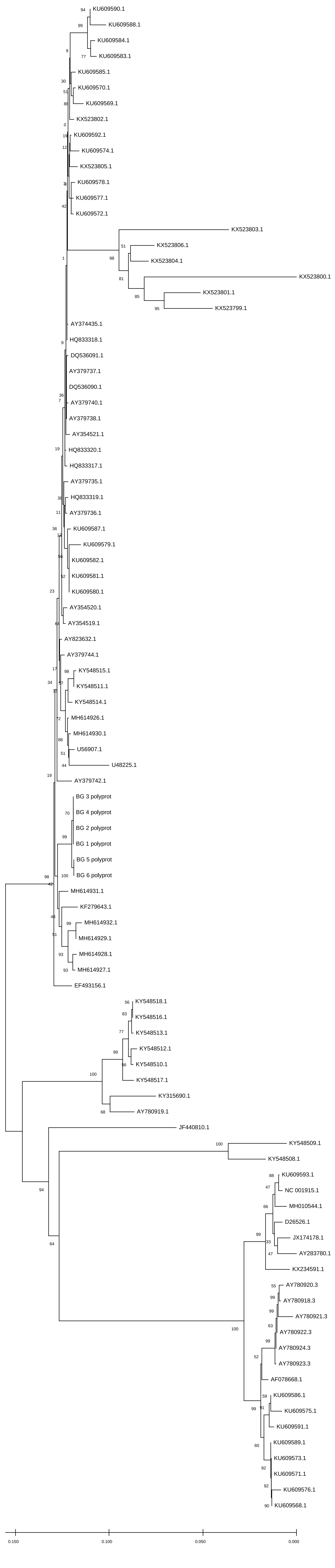

### Data Sheet 6.PDF

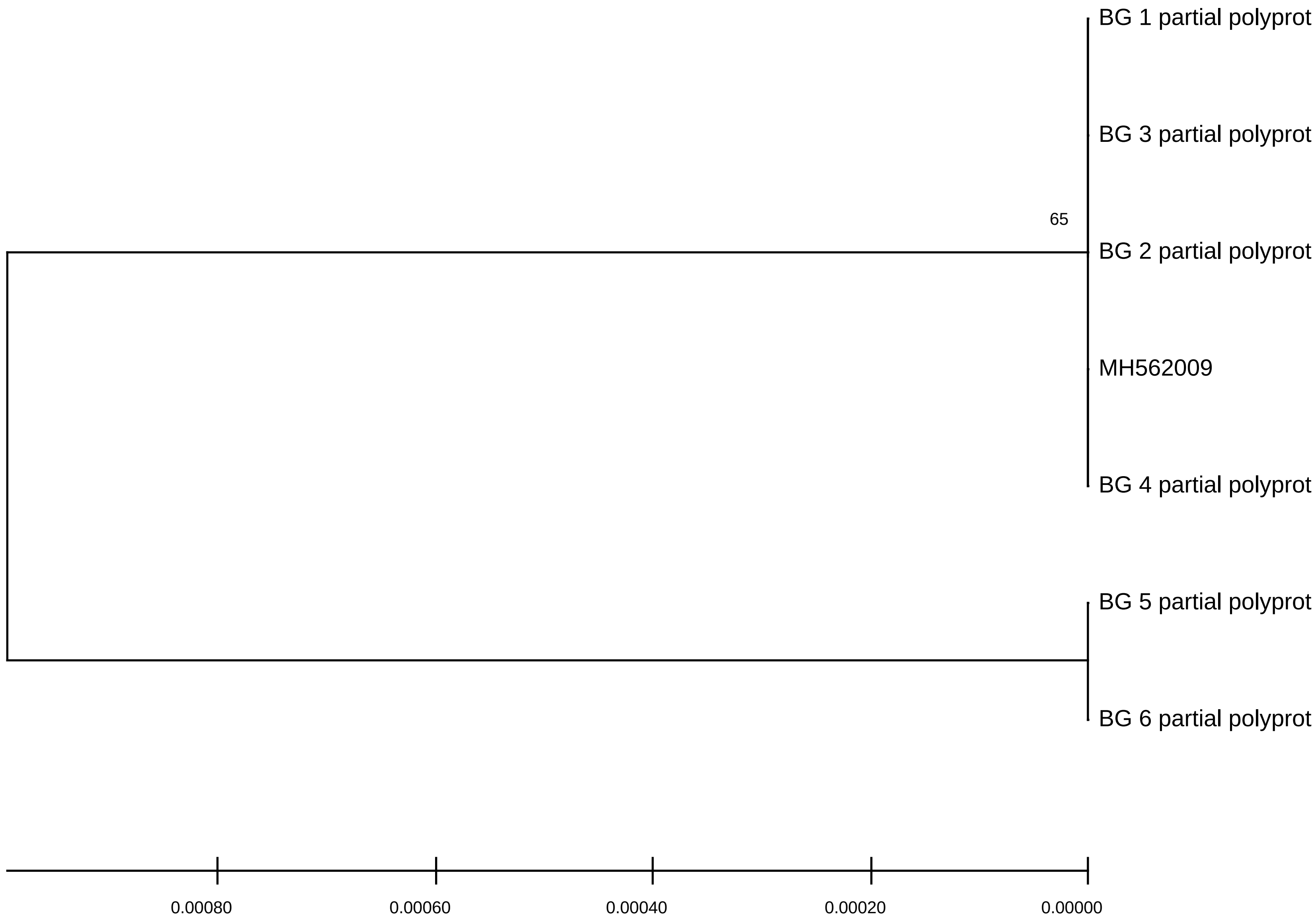

### Data Sheet 7.PDF

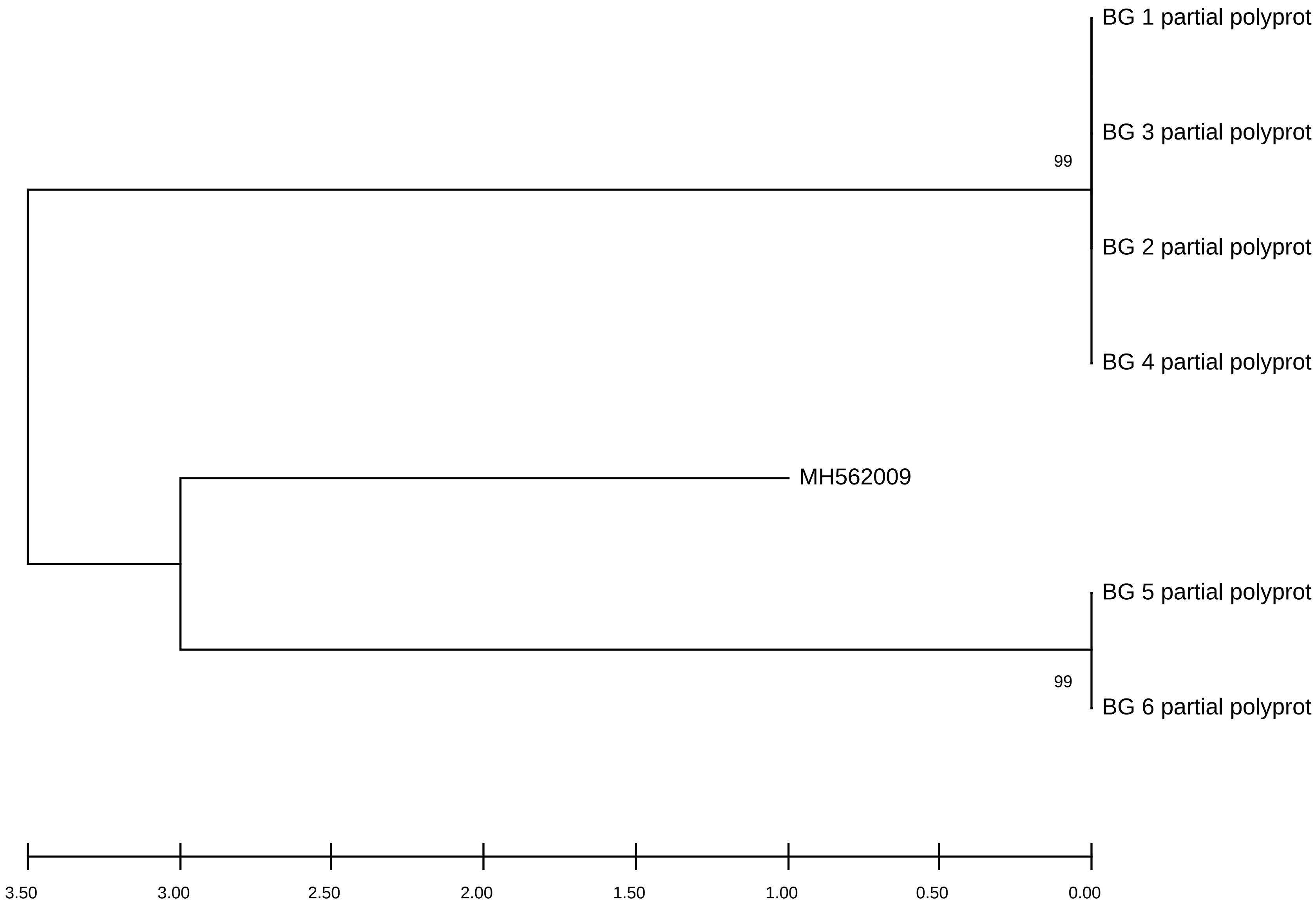
